# Extensions of the worm-like-chain model to tethered active filaments under tension

**DOI:** 10.1101/2020.07.26.222273

**Authors:** Xinyu Liao, Prashant K. Purohit, Arvind Gopinath

## Abstract

Intracellular elastic filaments such as microtubules are subject to thermal Brownian noise and active noise generated by molecular motors that convert chemical energy into mechanical work. Similarly, polymers in living fluids such as bacterial suspensions and swarms suffer bending deformations as they interact with single bacteria or with cell clusters. Often these filaments perform mechanical functions and interact with their networked environment through cross-links, or have other similar constraints placed on them. Here we examine the mechanical properties - under tension - of such constrained active filaments under canonical boundary conditions motivated by experiments. Fluctuations in the filament shape are a consequence of two types of random forces - thermal Brownian forces, and activity derived forces with specified time and space correlation functions. We derive force-extension relationships and expressions for the mean square deflections for tethered filaments under various boundary conditions including hinged and clamped constraints. The expressions for hinged-hinged boundary conditions are reminiscent of the worm-like-chain model and feature effective bending moduli and mode-dependent non-thermodynamic effective temperatures controlled by the imposed force and by the activity. Our results provide methods to estimate the activity by measurements of the force-extension relation of the filaments or their mean-square deflections which can be routinely performed using optical traps, tethered particle experiments, or other single molecule techniques.

## I. INTRODUCTION

The structural elasticity of slender biofilaments and polymers such as DNA, proteins, actin and microtubules and their deformation in response to their environment are key to their function^1–16^. In a homogeneous medium at constant temperature, embedded filaments and polymers suffer stochastic bending fluctuations. These deformations arise due to the action of thermal Brownian random forces on the filament. In passive settings, the energy to create a curved segment of the filament is drawn from the ambient bath; conversely, already curved segments of the filament flatten by viscous dissipation of stored energy into the medium. At equilibrium, this dual role of the thermal bath and the statistical properties of the forcing manifests in the fluctuation-dissipation theorem that relates the magnitudes of the delta-correlated forces with the thermodynamic temperature T^10,17–19^.

In this work, we study the properties of fluctuating elastic filaments under active settings. Such a situation can be realized in two ways - (Case A) the filament is surrounded by a suspension of active or living particles such as synthetic self-propelling colloids or living bacteria^20–25^, or (Case B) the filament is itself comprised of tightly and elastically coupled self-propelling monomeric agents^26^. As in the purely thermal non-active (passive) case, delta-correlated random forces are exerted as the bath molecules collide with the filament and deform it. In addition to these forces however, the filament is subject to athermal random forces due to interactions between the passive filament and the active particles as in Case A, or due to the random displacements between connected active particles as in Case B. In Case B, the active particles comprising the bead can change their orientation direction diffusively at intermediately long time scales independently of their translational motion. A related example in which the activity comes from the ambient is the mixed poroelastic active-passive matrix environment of eukaryotic cells^5,9–11,27,28^. Filaments in such settings can behave as described Case A; except that statistical properties of the activity and active stochastic forces (due to motor activity rather than free self-propelling bacteria) may be different^13^.

A key question in these problems is how local bending fluctuations, and indeed metrics for the size and shape of the filament as a whole, are related to intrinsic properties such as the coarse-grained effective bending modulus *K_b_* of the filament, its contour length *L*, the temperature of the ambient medium T and importantly the *activity*. An equally important question is to understand how these metrics impact the response of the filament to external stretching or compression forces - i.e, the force-extension relationship that encapsulates the coarse-grained elastic response of the filament - in the *presence of activity*. Finally, and external to the properties of the filament or of the embedding medium, is the role of constraints and boundary conditions. Filaments often perform mechanical functions and interact with their networked environment through cross-links, or have other similar constraints placed on them. Thus, boundary conditions are anticipated to play very important roles by constraining the space of possible configurations that can be attained.

Two powerful models have been traditionally used to study the stochastic deformations of filaments in passive settings - the Freely Jointed Chain (FJC) and the Worm Like Chain (WLC) models^10^. Of the two, the Worm Like Chain model (WLC) allows the incorporation of backbone elasticity as well as configurational entropy^1,2,4,5,7,29^ and is particularly suited for stiff to semiflexible polymers and biofilaments. This approach has been successful in describing the elastic properties of a variety of biomolecules, such as ssDNA, dsDNA, RNA and polypeptide chains (a comprehensive summary may be had from^10^, and bibliography therein). Consequently, WLC models and variants, appropriate for semi-flexible or stiff passive polymers/filaments have also been extended to active settings^30,31^; specifically properties of the filaments and complementarily, of the environment are determined in one of two ways.

The first approach is to study the fluctuations of freely suspended filaments. The geometrical size and shape of an ensemble of such suspended fluctuating filaments can be quantified by comparing *L* with effective radius of gyration *R_g_* and a suitably projected end-to-end distance *X*. Visual inspection of the ensemble of contour shapes provides the effective persistence length *ε* - the length scale over which parts of the full filament are orientationally correlated. Soft, long filaments such as DNA in water-like fluid at room temperature typically satisfy *ε* ≪ *L* and form blob-like compact conformations with *R_g_* ≪ *L*. Stiff filaments such as microtubules at room temperature do not adopt blob like configurations; their end to end distance ~ *L* and *ε* ~ *L*. In the absence of activity, and at equilibrium *K_b_* = *εk_B_*T where *k*_B_ is the Boltzmann constant. When thermal forces dominate elastic forces, as for a long soft blob-like polymer, the end-to-end distance ~ 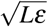; the radius of gyration scales similarly but with a different pre-factor. When activity is present, the relationship between *ε* and T is modified; bending energies corresponding to different Fourier modes are no longer related to T via the equipartition theorem. Here, activity serves as a source of energy but not as a sink - dissipation is ultimately due to passive viscosity effects. Recent papers on free active filaments or passive filaments in active fluids studied these novel properties in detail^30,31^.

Here, we focus on a second approach that deduces filament properties from a consideration of its quasi-static *stretching response to imposed pulling forces* under suitable chosen constraints. This approach is motivated by dramatic advances in experimental single molecule techniques such as atomic force microscopy (AFM), optical tweezers and optical traps, magnetic tweezers and pipette based force apparatus that allow stretching forces to be applied directly to the ends. These quasi-static experiments allow us to identify stiffening or softening responses as the polymer stretches by studying local variations of the force-extension curve as a function of extension.

Our starting point (Section II) is the Langevin equation mimicking a passive elastic curve bending under the influence of both thermal and active stochastic forces each with experimentally motivated stochastic properties and spatiotemporal correlations. The formalism follows Gaussian semiflexible polymer model proposed in Eisenstacken *et al*.^30,31^; similar models have been treated and studied elsewhere^32–34^. Active particles in this model move around in the constant temperature fluid with speed *V*_0_ and an effective viscosity dependent rotational drag coefficient *γ_R_*. Note that Eisenstacken *et al*.^30,31^ consider unconstrained suspended filaments with free boundary conditions. Here, we introduce the general solutions to the Langevin equations and obtain formal expressions for the mean square deformation and the mean stretch using appropriate eigenfunction expansions; these being controlled by the boundary conditions and nature of constraints. In Section III, we introduce three canonical boundary conditions motivated by experiments. General solutions to the Langevin equation are specialized to each of the three boundary conditions, and analytical (and in some cases, closed form) force-extension relationships and expressions are derived. The expressions for hinged-hinged boundary conditions are reminiscent of the worm-like-chain model and feature effective bending moduli and mode-dependent non-thermodynamic effective temperatures controlled by the imposed force and by the activity. In section IV, we analyze these expressions for realistic parameter values and illustrate how our results provide methods to estimate the activity from experiments. We conclude in Section V by motivating future avenues for theoretical and experimental work.

## II. ACTIVE POLYMER MODEL

### A. Langevin equation for active polymer

We will use the framework of Eisenstacken *et al*.^30,31^ in this paper but study the fluctuations of constrained active polymers relevant to biophysical and biological boundary conditions that, to the best of our knowledge, have not been studied elsewhere. In contrast to Eisenstacken *et al*.^30,31^ (a) we do not allow (at hydrodynamic time scales) motion of the center of mass of the filaments, so the filaments cannot translate but can explore the full range of bending fluctuations, (b) we work in a constant tension and constant temperature ensemble which allows us to deduce the force-extension relations for filaments immersed in active baths, and (c) we consider several boundary conditions that are applicable to different types of single molecule experiments. For example, the cantilever boundary condition considered in this paper can be applied to tethered particle experiments on filaments in active baths as a method to quantify the activity of the particles via deduction of *V*_0_ from routine measurements of bead fluctuations. In these constrained settings, we expect activity to further enhance differences between the WLC passive model and the active polymer model with non-thermal active fluctuations sometimes controlling filament configurations. Changes in the filament’s local curvature and curvature distributions are coupled to local injection and dissipation of energy by the actions of the active agents.

We start with the Gaussian semiflexible polymer model (or a fluctuating rod/filament model or an elastic line model)^30,31^ in which the polymer shape is completely defined by the locus of Lagrangian points coincident with the backbone at every time instant *t*. This in turn is treated as a continuous vector (curve) **r**(*t, s*), continuously differentiable, and parametrized by the arc-length parameter *s* with −*L*/2 < *s* < *L*/2. The evolution of **r**(*t, s*) is governed by a Langevin equation and has been derived earlier^29^.

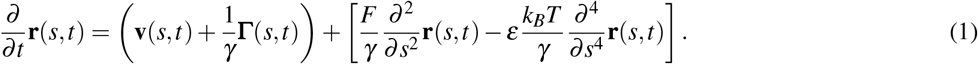

The meaning for each parameter in this equation is given in Table 1. Note that terms within parenthesis on the right are stochastic terms while the ones within the square bracket represents deterministic linear terms. In particular, *F* is the externally applied force assumed to be along the **e**_1_ direction in the lab frame. The decomposition **r**(*s, t*) = *x*(*s*)**e**_1_ + *y*(*s*)**e**_2_ provides components in two directions **e**_1_ and **e**_2_. Let *θ*(*s*) be the angle between tangent vector **T**(*t, s*) and **e**_1_ direction (see Fig. 1). The infinitesimal element along the filament *d***r**(*s, t*) = *ds***T**(*s, t*) and this element when projected along the two orthogonal axes provides the expressions *∂x* = *∂s* cos *θ* and *∂y* = *∂s* sin *θ*. Similar to the worm-like-chain model *F* is assumed to be large so that the deflections of the filament in the *y* direction are small enough that the angle in sin *θ* = *∂y*/*∂s* can be approximated as sin *θ* ≈ *θ* and *∂x*/*∂s* = cos *θ* ≈ 1 – *θ*^2^/2 (this is a good approximation in the range −*π*/6 ≤ *θ* ≤ *π*/6 as shown in Purohit *et al*^35^). In an ensemble where the end-to-end distance *X* is prescribed, *F* plays the role of a Lagrange multiplier and enforces the global constraint 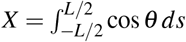. Here, we operate in an ensemble in which the force *F* is prescribed and the end-to-end projected distance *X* will be computed.

**FIG. 1.**
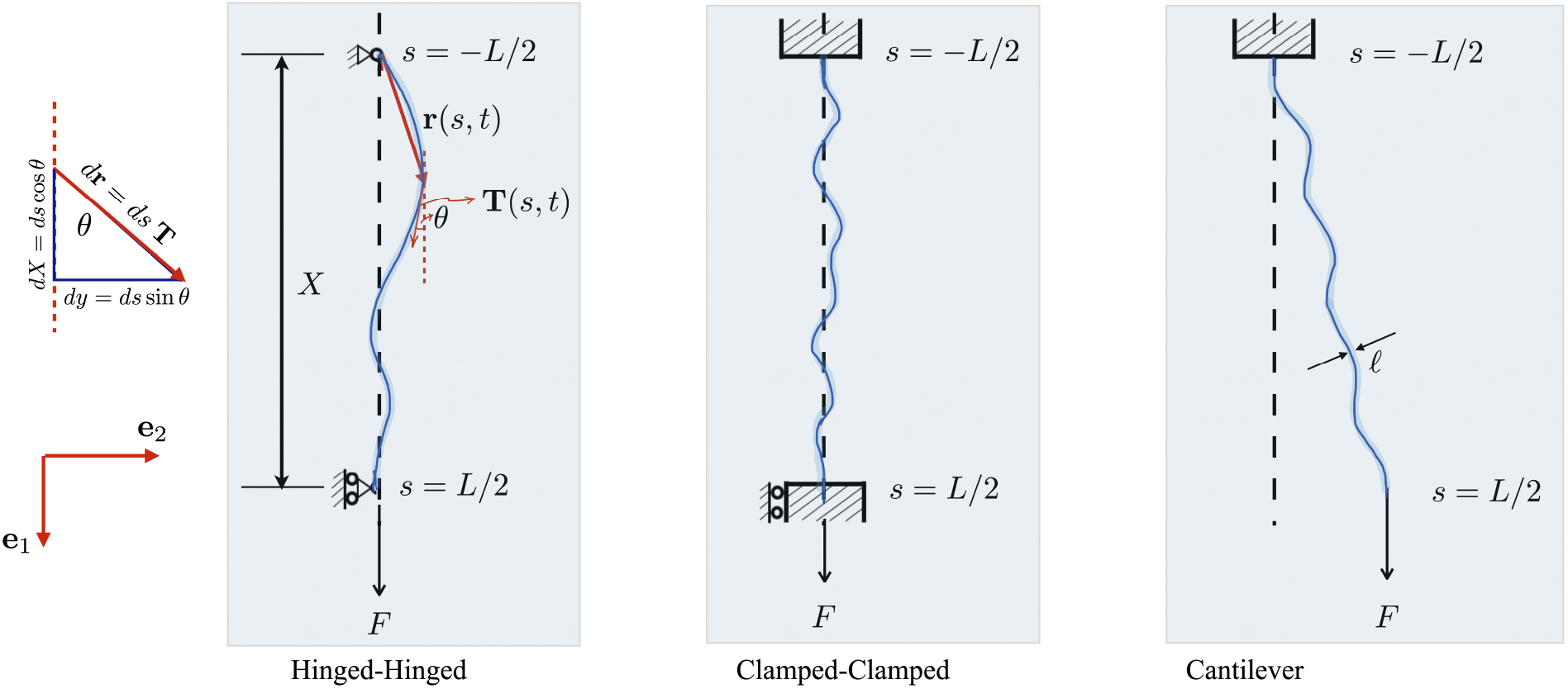
Schematic figures illustrating the three types of boundary conditions analyzed in this paper. In these scenarios, the active filament or polymer (blue curve) is immersed in a viscous medium. A force *F* is applied as shown in each case that causes the filament to stretch and orient in the direction of the force. The left-most tile shows projections along and orthogonal to the applied force *F*.

**TABLE I.** List of parameters

| Symbol | Description | Units | Typical values |
| --- | --- | --- | --- |
| $\ell$ | filament diameter | m | $7 \times 10^{-9}$ |
| $L$ | filament length | m | $6.82 \times 10^{-5}$ |
| $T$ | temperature | K | 300 |
| $k_B$ | Boltzmann constant | N·m·K<sup>-1</sup> | $1.38 \times 10^{-23}$ |
| $\gamma_R$ | damping factor | s<sup>-1</sup> | 0.16 |
| $\gamma$ | translational friction coefficient | N·s·m<sup>-2</sup> | $1.47 \times 10^{-3}$ |
| $F$ | applied tension | N | $1 \times 10^{-12}$ |
| $\varepsilon$ | persistence length | m | $1.71 \times 10^{-5}$ |
| $K_b$ | $\varepsilon k_B T$ | N·m<sup>2</sup> | $7 \times 10^{-26}$ |
| $V_0$ | propulsion velocity | m·s<sup>-1</sup> | $5 - 25 \times 10^{-6}$ |
| $\Gamma$ | stochasticity | N·m<sup>-1</sup> | $6.09 \times 10^{-24}$ |

Eqn. (1) exhibits the effects of two random processes - one thermal and one athermal (active). Following^30,31^, the term **Γ**(*s, t*) is a thermal component that arises from a stochastic Markovian, stationary and Gaussian process. This fluctuating force has zero mean and its second moments are given by

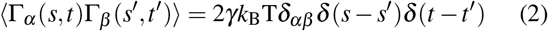

where T is the temperature, *k*_B_ the Boltzmann constant, and γ the translational friction coefficient per unit length. The influence of athermal noise originating in activity is captured by the velocity term **v**(s, t). This velocity is a non-Markovian but Gaussian stochastic process with zero mean and the correlation function^36^.

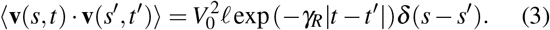

In reference to Case A, the velocity term is an approximation to the manner in which the active agents in the medium interact with the flexible passive filament (see also^37^ for the derivation of this equation). The ratio *L/ℓ* can in this case be interpreted^30,31^ as the number of uniformly distributed active sites along the polymer or equivalently as the number of possible sites active agents in the bath that can access and influence along the polymeric backbone. In reference to Case B, *V*_0_ is the self-propulsion speed of the active agent, *γ_R_* ~ *D_R_* its rotational diffusivity, and *ℓ* may be thought of as a characteristic thickness of the active agent.

We conclude this section by noting that eqn. (1) and subsequent analysis can also be used to analyze active motor-driven fluctuations of filaments. This is readily done by replacing the first term on the right involving the active velocity with a motor active force density using **v** = **F**^*m*^/*γ*. A simple approximation can be derived when motors acting along the filament are assumed to be in one of two states - either attached (on) or detached (off) - with attachment and detachment frequencies *ω*_on_ and *ω*_off_, respectively. The active force per motor 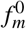 depends on the friction coefficient *γ* characterizing the ambient fluid properties as here, the duty ratio of the motor and details of the active force generating mechanisms. The active force density due to motors is then 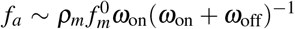 with *ρ_m_* being the line density of motors along the filament. Following previous studies^13^ we write

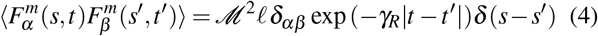

with 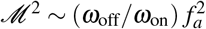 reflects the effective strength of the motor activity. Here *γ_R_* is to be interpreted as the inverse of an average time-scale for motor activity felt by the microtubule, *γ_R_* = (*ω*_on_ + *ω*_off_). With this mapping done, we go back to the form presented in eqn. (1)–(3) for analysis as below; however results presented in subsequent sections can be extended in a straightforward manner to scenarios where eqn. (1)&(2)&(4) are appropriate.

### B. WLC predictions for stretched passive polymers

Before we analyze the Langevin equation for the active filament/polymer further, we first summarize previous results obtained from WLC models for semi-stiff polymers in the absence of activity. For long WLC polymers of length *L* in thermal equilibrium, the persistence length *ε* and the mean square end-to-end distance are given by

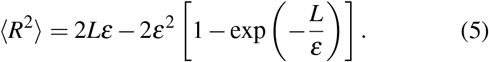

When such a polymer is stretched by applying tensile forces of magnitude *F* at the ends, the asymptotic expression for average end-to-end distance in the large force limit is 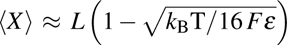. More generally, for short polymers or stiff polymers under tension and for large forces,

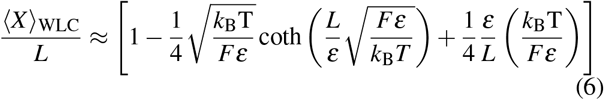

provides an experimentally useable form of the force-extension relationship^35^. For arbitrary cases, the more accurate form^12^

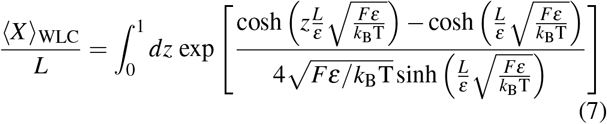

have been used. As a point of comparison, we consider that the force-extension relationship in a purely entropic framework using the FJC model. Let the FJC polymer of length *L* be comprised of *N* straight rigid freely jointed (Kuhn) segments each of size *ε*. The mean projected length of the polymer when held fixed at one end and stretched by a force *F* at the other is

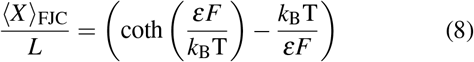

Eqn. (5)–(8) prominently feature the ratio of two energy scales, the force derived scale *Fℓ*_c_ and the thermal scale *k*_B_T. For the WLC model, the relevant length scale *ℓ*_c_ = *ε*. These expressions serve as a starting point to next look at the effects of activity.

### C. Scaled parameters quantifying activity and elasticity

Based on eqn. (1)–(3), we now introduce important dimensionless parameters that play a role in subsequent analysis.

We first consider activity independent quantities. Three independent aspects here need to be taken into account - the geometry of the filament, the elasticity of the filament based on persistence lengths at the thermodynamic temperature T, and the effect of the imposed force *F*. The dimensionless parameter 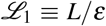 with *ε* ≡ *K_b_*/(*k*_B_T) being the thermal persistence length quantifies the effective stiffness at temperature T. When *L* » *ε*, the filament in the absence of activity behaves as a Gaussian chain. A second dimensionless quantity is 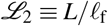 where the length-scale for the persistence of force 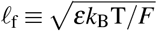. This length-scale determines the extent to which thermal fluctuations locally reorganize and randomize alignment along the polymer. As the force *F* increases, *ℓ*_f_ decreases, and for fixed *L* so does 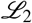 indicating that the filament is increasingly oriented with the direction of *F*.

It is also useful to define a dimensionless force

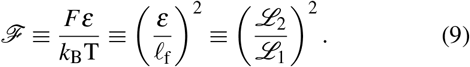

When 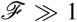, the polymer backbone is stretched predominantly along the direction of the force and thus the local tangent angle measured with respect to the direction of *F* is nearly zero all along the filament. Note that *F* → 0 with geometric and material properties fixed implies *L/ℓ*_f_ → 0 and 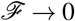. Setting *K_b_* → ∞ implies *ℓ*_f_ → ∞ and *L/ℓ*_f_ → 0.

Finally, we introduce parameters that incorporate the characteristics of the active agents. A third dimensionless scale, 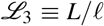 may be defined that provides an estimate for the number density of activity agents or sites. Additionally arising from the self-propelling nature and persistence times associated with the active agents (eqn. (2)&(3)), and matching previous studies, we define a Peclet number^30,31^

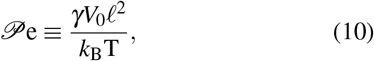

For unconstrained filaments, 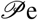 is important since the center of mass of the active filament can move and given the linearity of eqn. (1) we expect terms of the first power in *V*_0_ to control this motion. For constrained filaments, fluctuations in orientation will be important and this suggests that a different activity parameter will characterize the dynamics.

### D. Eigenfunction expansions deliver general expressions for mean square displacements and mean stretch

Following Eisentstacken *et al*.^30,31^, we assume that **r**(*s, t*) has an eigenfunction expansion given below where *φ*(*s*) are the orthonormal eigenfunctions of the deterministic and linear part of the stochastic differential equation. Filaments can generally move in a plane or in 3-D. Here we consider the planar motion of the filament and decompose the position vector in terms of orthonormal basis functions

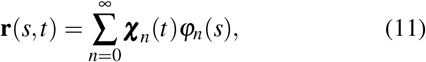

which yields the equation for the projection *y*(*s, t*)

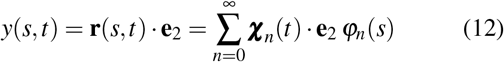

with the associated orthonormality condition on the eigenfunctions *φ_n_*(*s*):

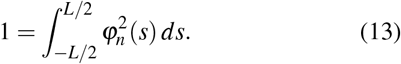

Substituting eqn. (11) in eqn. (1), taking projections and making use of the orthogonality of the eigenfunctions, we find that *φ_n_*(*s*) satisfies

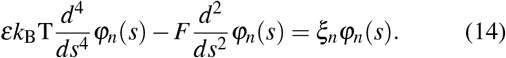

The general solution of eqn. (14) is given by,

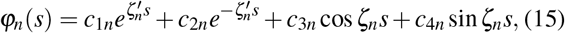

where

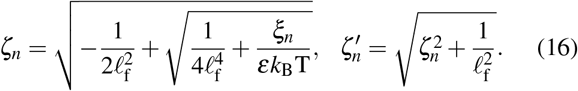

The mode amplitudes in eqn. (12) are related to the stochastic forcing which are the origin of time dependence^38^. Following the original derivation^30,31^ and motivated by the linearity of eqn. (1), we look for stationary functions by decomposing the fluctuating stochastic terms in eqn. (1) in terms of the same eigenfunctions as in eqn. (12). Thus we write,

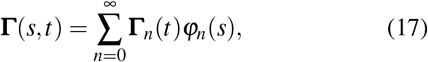

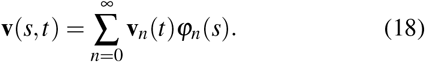

Substituting eqn. (17) and eqn. (12) into eqn. (1), we reproduce the primary result from^30,31^

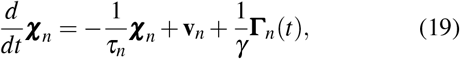

with

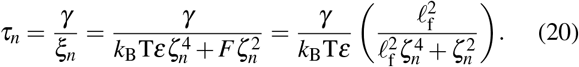

To further proceed and calculate properties such as the end-to-end stretch we need analytical results for the mode amplitudes. This has been derived earlier in^30^ and we report them here for completeness. The solution and associated orthogonality conditions are

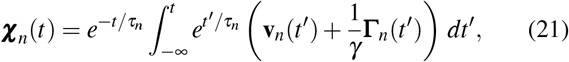

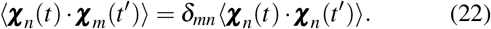

Time correlations have been calculated elsewhere (see eqn. (23) in^31^). In eqn. (22) *δ_mn_* is the Kronecker delta function. In the rest of the paper we will confine attention of the component of **r**(*t, s*) along the **e**_2_ direction, thus we will only be interested in the **e**_2_ component of ***χ**_n_*. This component will be denoted by the scalar *χ_n_*. From complete expressions in^31^, we derive

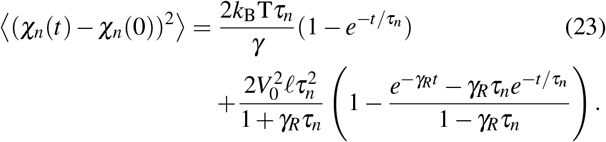

The statistically stationary properties are obtained in the limit where the memory of initial conditions is lost; thus taking the limit *t* → ∞ in eqn. (21), we can calculate ensemble averaged correlation functions in the long time limit. Further insight may be had by examining in certain limits. The small time expansion of eqn. (24) confirms that

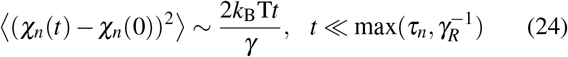

and independent of *F*. We note that *τ_n_* depends on *F* via eqn. (20). When 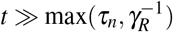

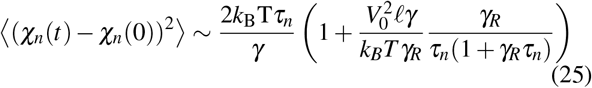

and thus strongly dependent on both *F* and *V*_0_.

### E. Expressions for the mean square displacement

Eisenstacken *et al*.^30,31^ considered free filaments that are allowed to diffuse and therefore have a non-zero mean square displacement (stationary state value), that is the 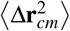 is in general non-zero. Here, we consider constrained filaments that are prevented from global translation and therefore ignore this term.

The mean square value of the *y* displacement *y*(*s, t*) with −*L*/2 ≤ *s* ≤ *L*/2 at steady state can be obtained by taking the statistically stationary average as *t* → ∞ of eqn. (24) which gives

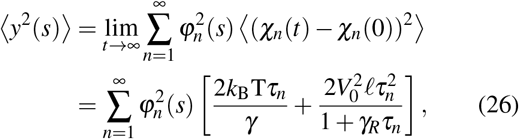

where eqn. (24) has been used in going from the first to the second expression on the right hand side. We note that activity enters through the eigenfunction as well as the multiplying factor in square brackets while the tension ~ *F* enters explicitly through the eigenfunction.

Note that the two terms in the square bracket in eqn. (26) provide information about two complementary features of the activity. The first term gives an indication of the time scale over which correlations due to thermal noise decay (via *τ_n_*) and this incorporates the effect of *F* (see eqn. (20)) but does not contain the influence of activity. The second term on the other hand explicitly incorporates the effect of active self-propulsion from its dependence on *V*_0_ and also the geometry of the agents through *ℓ*.

### F. Expressions for the mean stretch

Apart from mean square displacement, we are also interested in the force-extension relation. We use *X* to denote the projected length of the filaments onto the *x*-axis (see Fig. 1), so that *dX/ds* = cos *θ*(*s, t*) where *θ*(*s*) is the angle between unit tangent vector *d***r**/*ds* at point *s* and the **e**_1_-axis. We assume that the displacements are small and sin *θ* ≈ *θ* so that

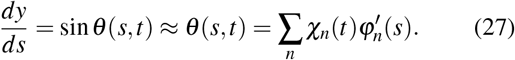

Next, using cos *θ* ≈ 1 – *θ*^2^/2 we obtain

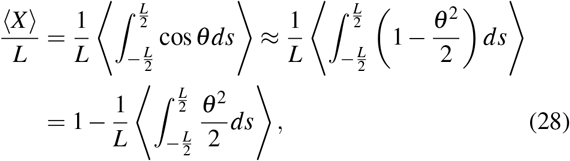

which evaluates to

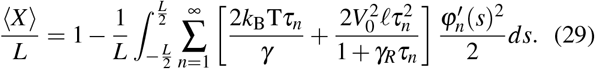

## III. RESULTS: ANALYSIS OF CONSTRAINED ACTIVE FILAMENTS

Here, we begin with the general forms of the equations in the previous section for the eigenfunctions given by eqn. (14)(16), equations for the mode amplitudes given by eqn. (17)(21) and the analytical expressions eqn. (26)&(29). Specific forms for the eigenfunctions are determined for different boundary conditions, and from these reduced expressions for eqn. (26)&(29) for the case at hand is derived and analyzed.

### A. Hinged-Hinged boundary condition

Under hinged-hinged boundary conditions, the moments at the ends of the filament are zero and this is usually modeled in mechanics by hinges which allow free rotation as illustrated in Fig. 1. The boundary constraint is therefore soft in that the hinges resist forces and prevent translation degrees of freedom but allow for free rotation. The active filament is subject to an applied tension acting in the *x*-direction at one end as in the figure. The hinge at the end where the force is applied is on a roller so that it can move in the *x*-direction to change the end-to-end distance, or effective extension of the filament.

Note that the hinged ends can support forces. Specifically, the roller does not allow motion in the *y*-direction (perpendicular to the tension), hence there is a reaction (shear) force at the end. This reaction force does not enter the potential energy of the filament since it does not do work. Experimentally, the hinged-hinged boundary condition may be realized by using an optical trap (which allows free rotation) to pull one end of the filament while the other end is held in a second optical trap or is immobilized by some other means.

The appropriate boundary conditions that need to be imposed on eqn. (12) in this case correspond to the constraints of no displacement and zero moment. The curvature *κ* = |*d*^2^**r**/*ds*^2^| which can be approximated by *κ* = *∂*^2^*y*/*∂s*^2^ for small *y*. Assuming that the moment is proportional to *κ* as in classical Kirchhoff-Love rod theory, the boundary conditions can be written as

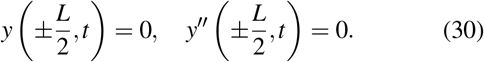

where the ′ denotes differentiation with respect to *s*. Substituting eqn. (30) into eqn. (15), and imposing that the *s* dependence of *y*(*t, s*) arises only from *φ*(*s*) in the eigenfunction expansion eqn. (12), we obtain the eigenvalues and eigenfunctions (see Appendix A)

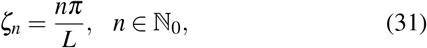

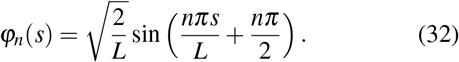

Thus, the eigenvalues and eigenfunctions take a particularly simple form that is amenable to analysis using Fourier series. With these, we can now evaluate statistical properties of this constrained elastic filament.

#### 1. Mean square displacement

Eqn. (26) can be rewritten (eqn. (A1)–(A8)),

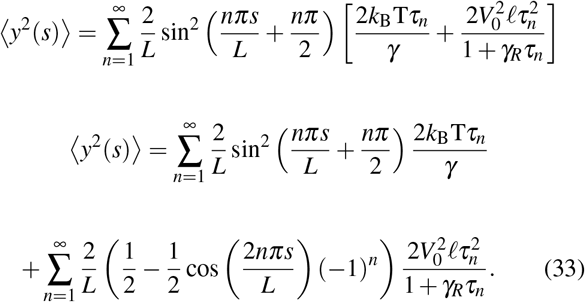

To ease the analysis, we further introduce intermediate variables

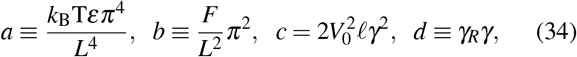

and

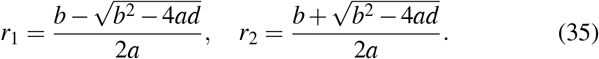

Simplifying eqn. (33) further (see eqn. (A9)–(A13)), and using eqn. (34)&(35) to rewrite the results, we obtain the general expression (eqn. (36)) and the expansion for large *F* (eqn. (37)) as below:

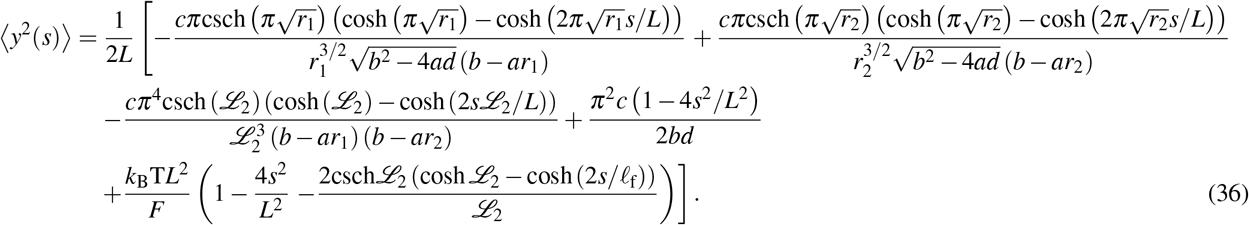

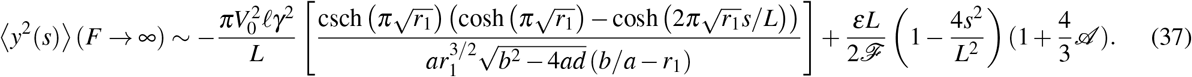

The last term of eqn. (37), introduces a dimensionless quantity that represents the ratio of energy scales - an effective active energy to the thermal energy, *k*_B_T:

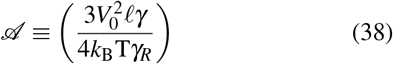

Here we note that 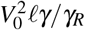 is the product of the active force *V*_0_*γℓ*, and the characteristic distance over which this force persists *ℓ*(*V*_0_/*γ_R_*). Thus 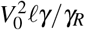 is a measure of the amplitudes of the fluctuating active energy deforming that filament.

#### 2. Force-extension relation

The force-extension relation for a hinged-hinged filament in an active bath can be derived from eqn. (29). We start with the equation for the statistically averaged mean stretch that is first written as an infinite series in *n* as in eqn. (33) below and then manipulated using steps similar to that described above. We note that the Langevin equation is linear in the activity (specifically in 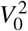). Algebraic manipulations allows us to go from eqn. (33) to the closed form expression

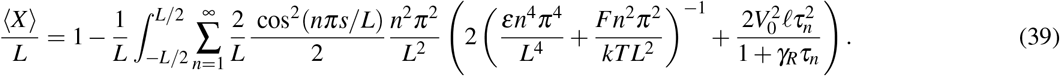

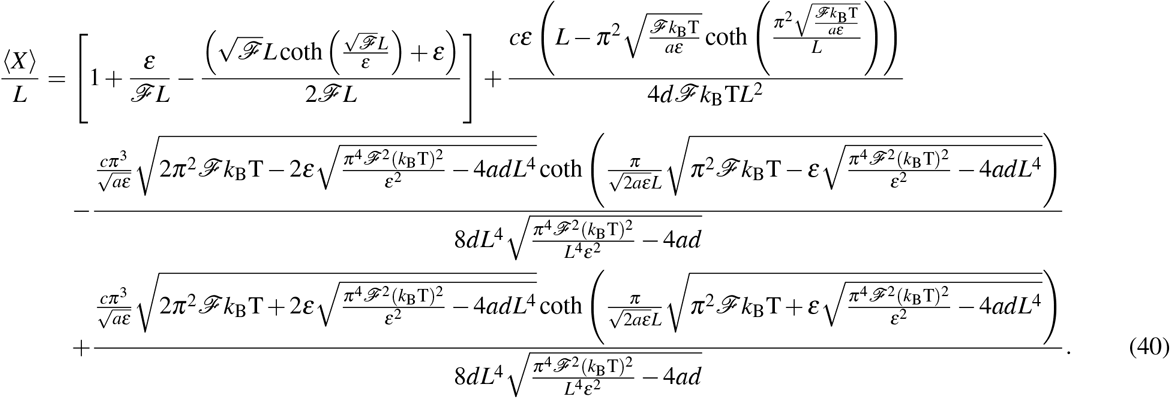

The terms within the square brackets on the right hand side are independent of activity; other terms on the right hand side depend on *c* linearly.

As 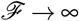, we can ignore 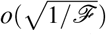 in eqn. (40) and simplify further to obtain the mean stretch for large force *F* and arbitrary values of 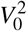 (arbitrary values of *c*):

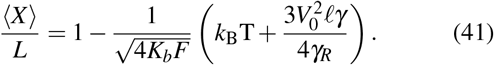

The Worm-like chain formula can be recovered from eqn. (41) when *V*_0_ = 0, which is as expected. Eqn. (41) demonstrates that, as far as the mean stretch is concerned, in the large *F* limit, the filament behaves as if it is suspended in a continuous medium with an enhanced higher temperature given by

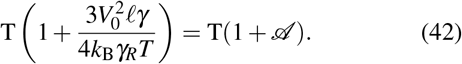

### B. Modes fluctuate at *effective* higher temperatures

Several studies on the thermodynamics of active suspensions have introduced the concept of an *effective temperature* that takes into account the enhanced effect of activity induced fluctuations. For instance we have recently studied^23,25,39^ if such a quantity can be defined in a dilute bacterial suspension of *Escherichia coli*^24^ by studying the diffusivity of small tracer particles and their speed distribution functions. We found that while useful in interpreting the enhancement in tracer diffusivity, the temperature is not a thermodynamic quantity^23,25^. Effective temperature concepts have been used to study various aspects of active biological matter such as red-blood-cell membrane fluctuations^28^, and active matter in general^28,40^, amongst others (c.f discussion in^25^). The equipartititon theorem applied to passive fluid interface^41,42^ and active bacterial interface fluctuations^43^, which like deformation modes in this study store energy, similarly provide a relationship between deformation modes and the thermodynamic temperature of the system. This has been used to relate properties such as surface tension and surface energy to the temperature.

Brangwynne *et al*.^27^ analyze the fluctuations of a microtubule which is embedded in a network of actin filaments. When no myosin motors are put into the actin gel then the microtubule suffers purely thermal fluctuations as seen in Figure 2A of their paper. When myosin motors and ATP are added into the mix then the microtubule is in an active environment and so the amplitude of its fluctuations increases as seen in Figure 2C of their paper. Here we use the eigenvalues and eigenmodes we have calculated in Section III to see if one can interpret the fluctuations seen in their experiments (Figure 2C in their paper) as due to an enhanced effective temperature.

**FIG. 2.**
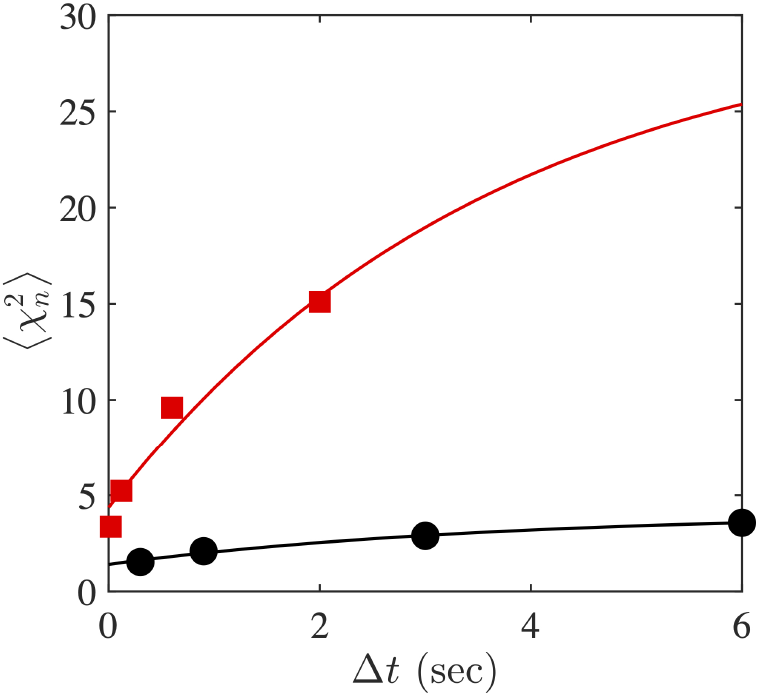
Black data and line correspond to *q* = 0.7mm^−1^ from Figure 2A of Brangwynne *et al*.^27^, red to *q* = 0.7mm^−1^ from Figure 2C of the same paper.

To proceed in our analysis, we revisit eqn. (26) for transverse fluctuations of the active filament given by eqn. (1)–(3) describing statistics of the random stochastic terms. Let *y*^2^(*s*) denote

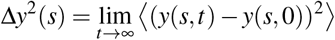

where without loss of generality, we assume that information about the initial shape is lost eventually. Noting that *τ_n_* is independent of activity, we rewrite eqn. (26) as follows

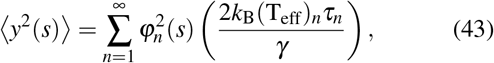

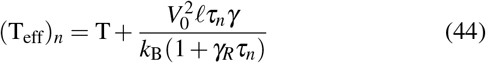

where we have introduced a mode dependent *effective temperature* (T_*eff*_)_*n*_ that differs from the thermal temperature (of the solvent) T when 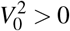. Eqn. (44) indicates that each mode *n* differs in dynamics through the factor *ζ_n_*. The actual form of the effective temperature will depend on the force exerted; this is due to the presence of *τ_n_* in the activity term - the second term on the right hand side of eqn. (44).

From the caption of Figure 2 in^27^ we find *L* = 30 mm and from the text *ε* = *K_b_*/*k*_B_T = 1 mm and thus 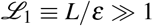; in this limit, and for negligible deterministic forces acting along the filament (as in their problem) the boundary condition does not matter in the computation of the amplitudes of the normal modes. Consideration of the basis functions used in their paper suggests that the microtubule ends are free to rotate being torque free and hence a hinged-hinged condition is appropriate. Consistent with their expression, and using eqn. (31)&(32) in eqn. (43) we get equations for the tangent angle and the curvature 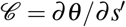

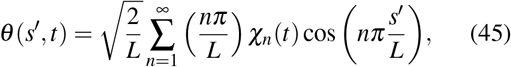

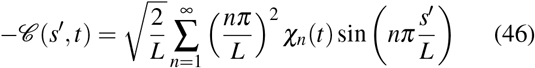

where we have transformed from used *s*′/*L* = *s/L* + 1/2 to transform from our range *s* ∈ (−*L*/2, *L*/2) to their *s*′ ∈ (0, *L*). Choosing the base preferred curvature to be zero, the ensemble averaged bending energy for *F* = 0 is

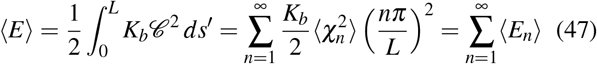

where we have exploited orthonormality of basis functions.

In the *absence of activity*, equipartition holds and thus the average energy in each mode of fluctuation per degree of freedom is 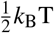. In two dimensions, equating each mode 〈*E_n_*〉 to *k*_B_T we obtain,

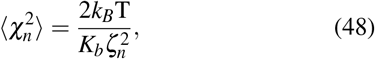

which is the expected result for *V*_0_ = 0 and reproduces the result in Brangwynne *et. al*.

In the presence of motors, the system is not thermodynamic anymore, and activity causes fluctuation amplitudes to increase relative to the thermal amplitudes. We then ask if an extended formula

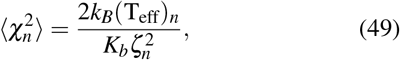

may be used to interpret experimental measurements of microtubule fluctuations in the presence of motors added. Note that the form of eqn. (44) emphasizes how activity enables this system to break free of some of the constraints imposed by fluctuation dissipation - specifically, activity leads to enhanced amplitudes, especially at small mode numbers *ζ_n_* due to the second term being a function of *τ_n_*. Note that we do not have detailed knowledge about motor activity (such as the motor forces, duty ratios, and attachment probabilities) needed to quantify the parameters in eqn. (4).

Eqn. (24) indicates that when activity is absent, 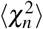 is approached exponentially in time *t*. Figures 2A (and 2C) of Brangwynne *et al*.^27^ give 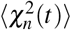 at four different values of *t*. Choosing *ζ_n_* = 0.7mm^−1^ and fitting the four values in time from Figure 2A to a simple decaying exponential

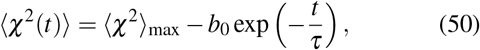

we find *τ* = 3.63 seconds and *b*_0_ = 2.67 nm consistent with *k*_B_*T* ≈ 4.1pNnm. The data points and the fit are shown in black in Figure 2.

When activity is present eqn. (24) suggests a trajectory in time that is no longer a simple decaying exponential; indeed setting *F* = 0 in eqn. (24) shows the presence of a constant time independent term and two decaying exponentials with opposite signs. For simplicity, here we however retain the form in eqn. (50) and treat 〈*χ*^2^〉_max_, *τ* and *b*_0_ as functions of activity. Choosing the same value of *ζ_n_*, we fit the data extracted in Figure 2C, and obtain *τ* = 3.63 seconds, *b*_0_ = 26 nm, and *k_B_*T_eff_ ≈ 30.5pNnm suggesting a significant activity driven component to the microtubule fluctuations. These points and the fit are plotted in red in Figure 2. These curves are to be compared with Figure 3 of^27^ in which the curve for 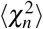 also flattens out as *t* becomes large.

**FIG. 3.**
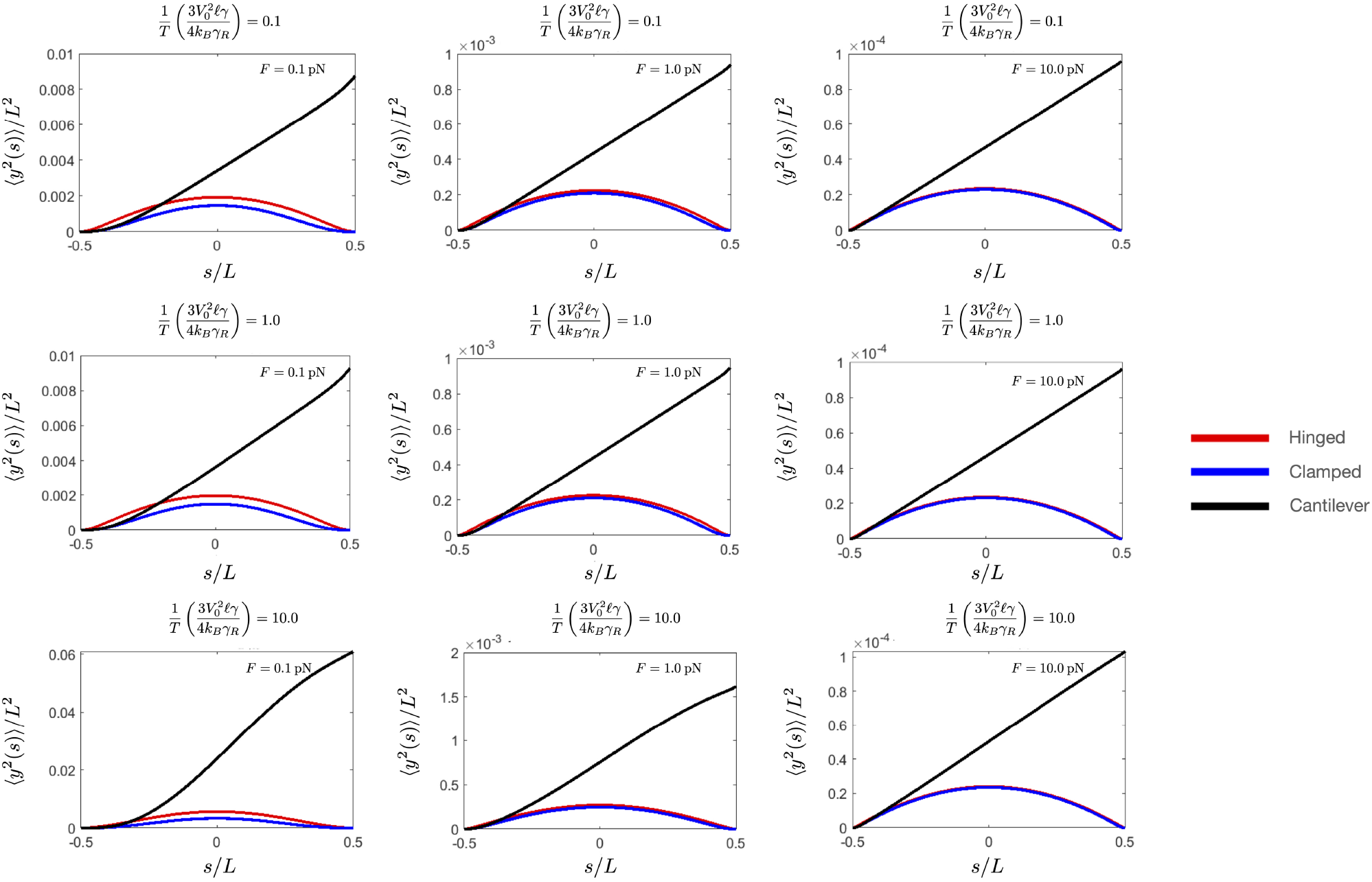
Mean square displacement under three types of boundary conditions. *ε* = 17.1 *μ*m, *L* = 8.54 *μ*m, *V*_0_ = 9.22 *μ*m/s, *ℓ* = 7 nm, T = 300 K, *γ_R_* = 0.16/s, *γ* = 1.47 × 10^−3^ N·s/m^2^.

**FIG. 4.**
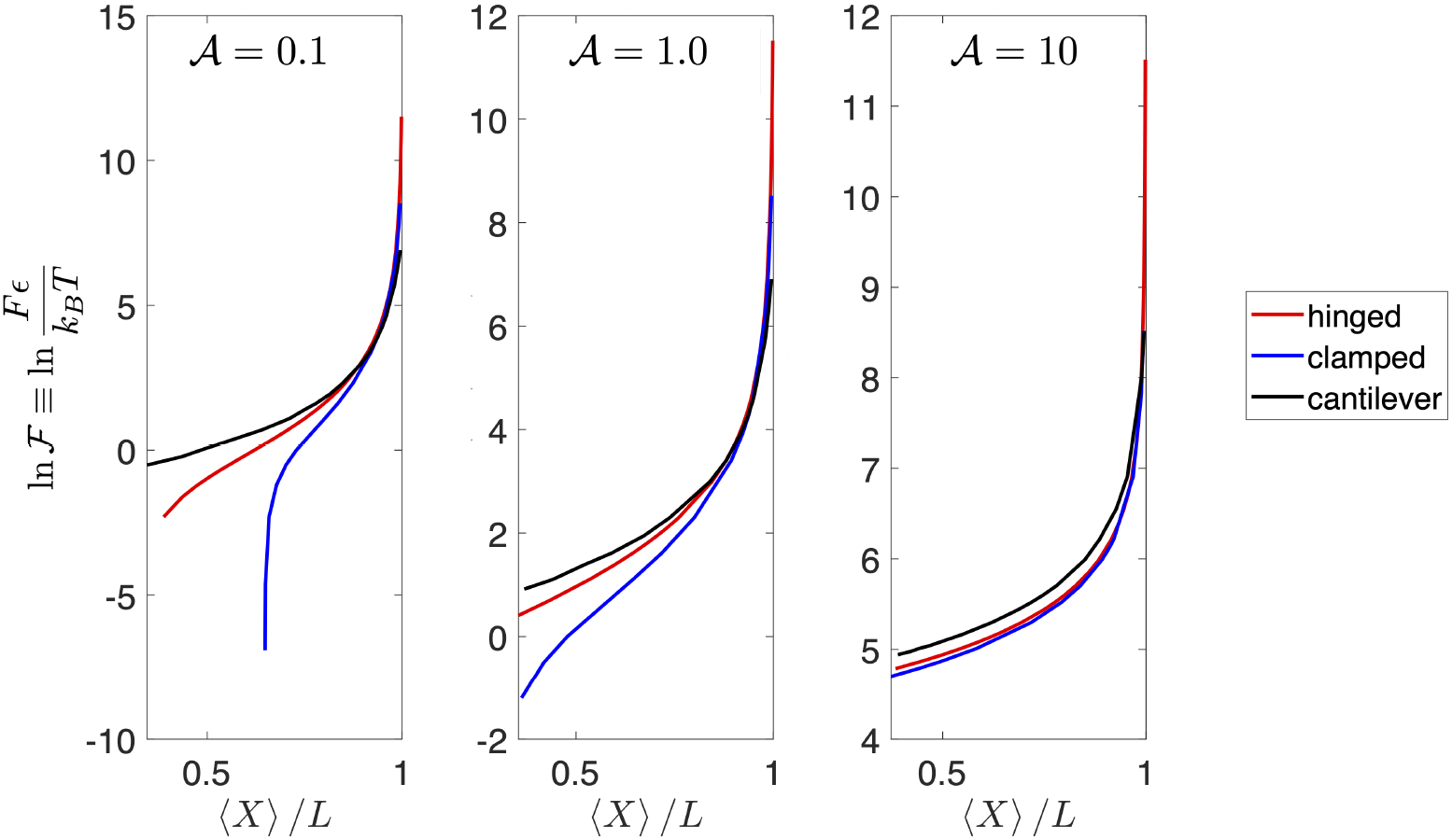
Force extension relations under three types of boundary conditions. The filament length used there is *L* = 68.2*μ*m. Other parameters are the same as Fig. 3.

### C. Clamped-Clamped boundary condition

#### 1. Mean square displacement

Clamped-clamped boundary conditions are relevant when no *y* displacements or rotations are allowed at both ends of the filament. Thus, there are non-zero shear and moment reactions at the two ends, but they do not enter the potential energy of the filament because they do no work. This type of boundary condition is relevant in assays in which filaments (actin or microtubules) bridge a channel (see Arsenault *et al*.^44^) while both their ends are attached to the substrate. This type of assays are often used to study the motion of molecular motors as they walk along the filament (the channel allows the motors to rotate around the filament without hindrance). The filaments can be under tension before being attached to the substrate as Arsenault *et al*.^44^ demonstrated by using electric fields in the case of actin filaments. It is not difficult to visualize a single molecule experiment in which a clamped-clamped boundary condition is applied together with a tensile force at one movable (in the *x* direction) end.

The clamped constraint at each end means that the angle made by the filament with the **e**_1_ axis is fixed. Here we assume that the clamping keeps the filament perfectly aligned with the direction of the force at *s* = ±*L*/2. The boundary conditions for the clamped-clamped filament are:

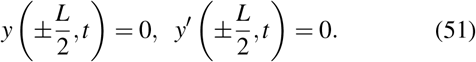

Dragging eqn. (51) into eqn. (15) yields the following two sets of solutions (see Appendix B). The first eigenvalue eigenfunction set is

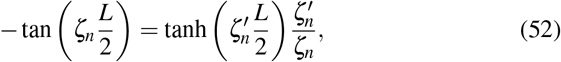

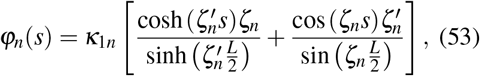

where 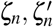 can be solved from eqn. (16)&(52). The second set is given by the two equations

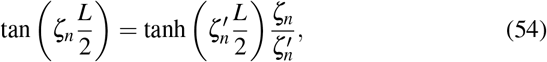

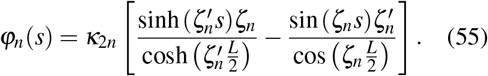

Here, *κ*_1*n*_ and *κ*_2*n*_ are given by,

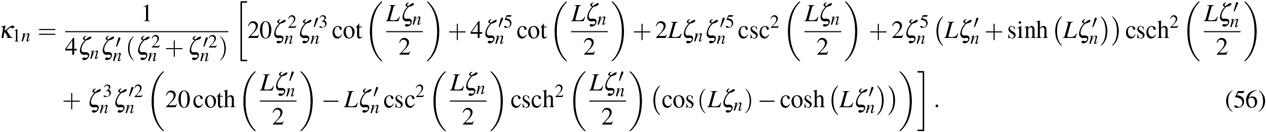

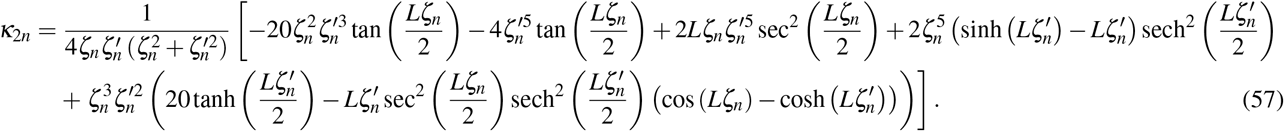

Lacking any straightforward algebraic manipulations that can yield a closed form expression as in the hinged-hinged case, we just use the series summation to evaluate the mean square displacement. Using 100 terms in the series proved more than adequate; 10 terms were insufficient indicating that the series, while converging to the exact value, does so moderately fast.

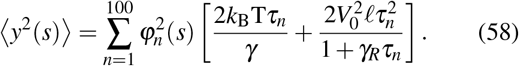

#### 2. Force-extension relation

There is no closed form solution for eqn. (29) under clamped-clamped boundary condition, so here as well we sum the first *N* = 100 terms of the series to obtain a sufficiently accurate value (here we use eqn. (53)&(55) to compute 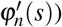.

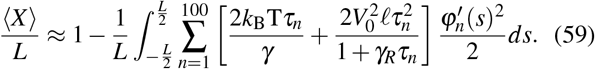

### D. Cantilever boundary condition

The cantilever boundary condition corresponds to one free end and one clamped end. The free end has zero shear force and moment, while the clamped end has both shear and moment reactions that do no work. This boundary condition is relevant to tethered particle experiments in single molecule mechanics. Often, the deflections of the tethered particle are measured in these experiments to deduce the length of the tether. A tensile force can be exerted on the particle by the use of magnetic tweezers. This tensile force does work through the motion of the bead and thus changes the end-to-end distance of the filament.

#### 1. Mean square displacement

We next calculate the mean-square displacement by first finding the associated eigenvalues and eigenfunctions.

The cantilever boundary conditions are,

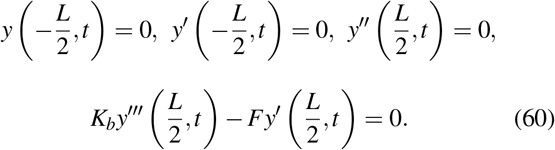

Re-defining 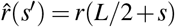, we rewrite the general solutior for eigenfunction eqn. (15) in the following way,

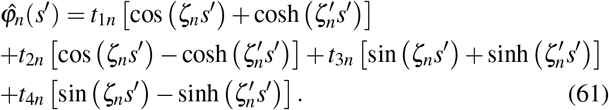

The first boundary condition 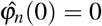 implies *t*_1*n*_ = 0. The second boundary condition 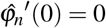 implies,

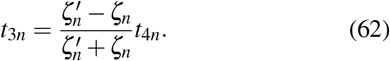

Using eqn. (62), the third boundary condition 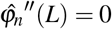 implies

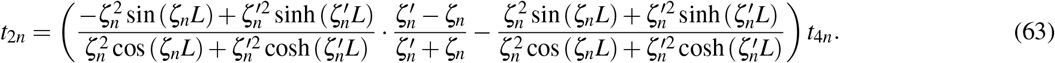

Imposing the boundary condition 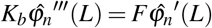 (to get eqn. (64)) and combining eqn. (62)–(64), we get eqn. (65),

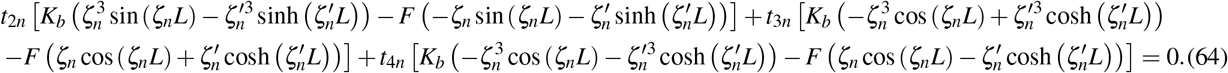

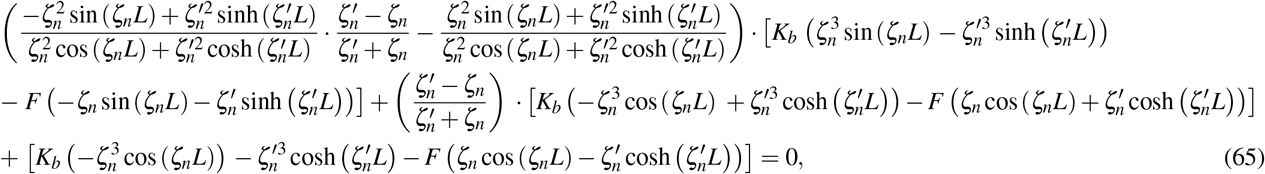

where the wave numbers *ζ_n_* and 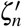 can be solved from eqn. (16)&(65). Using 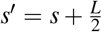, the eigenfunction can be written down in closed form

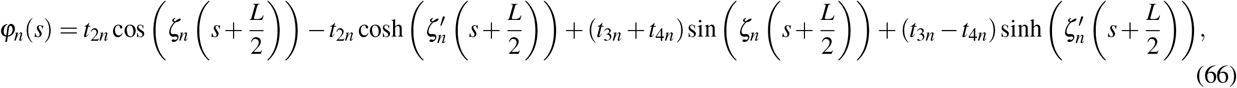

where the constants *t*_2*n*_ and *t*_3*n*_ are functions of *t*_4*n*_ through eqn. (63)&(62). The constant *t*_4*n*_ is solved by imposing the constraint 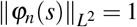 and orthonormalizing the eigenfunctions using eqn. (13). Similarly, we used eqn. (58) to compute 〈*y*^2^(*s*)〉.

#### 2. Force-extension relation

Again, there is no closed form solution for eqn. (29) under cantilever boundary condition, so we use eqn. (59) to approximate the force extension relation and eqn. (66) as the appropriate eigenfunctions corresponding to the cantilever boundary condition.

## IV. DISCUSSION

Here we analyze the expressions derived earlier and study how active noise modifies statistical properties for biologically realistic values of the tension *F*. In section III, we started from the evolution of polymer configurations using an *active extension* of the classical Langevin equation that has two stochastic components - the active noise due to the self propulsion velocity **v**(*s,t*) and the thermal fluctuating drag **Γ**(*s,t*). We then obtained explicit expressions for statistical stationary state averages for the mean square displacement and for the mean stretch of an active polymer subject to a tension in an ensemble where the temperature T is held constant. Contrary to other related works, the analysis in this section was specifically for tethered not free polymers. Associated deformations and effective force-extension relationships therefore depend crucially on the type of constraint at the boundary.

### A. Predictions of theory for weak and strong tension

We choose the tension *F* applied at *s* = *L*/2 to have values 0.1 pN, 1 pN, and 10 pN. The appropriate parameter to study in order to elucidate the effects of activity is the dimensionless parameter 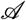. We choose 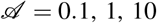 spanning low activity, moderate activity and high activity regimes. Note that since the geometric properties of the filament and the thermal temperature T is held constant, activity variations correspond to changes in the self-propulsion speed *V*_0_.

We compare the mean square displacement for the three types of boundary conditions under these parameters - these are given by eqn. (30)&(51)&(60) respectively. The results are plotted in Fig. 3. We see that 〈*y*^2^(*s*)〉 is maximum at *s* = 0, or the center of the filament, for the hinged-hinged and clamped-clamped boundary conditions. For the cantilever boundary condition 〈*y*^2^(*s*)〉 is maximum at the end *s* = *L*/2 as expected. The mean square displacement is predicted to be the smallest for the clamped-clamped filaments because they are the most constrained. From Fig. 3 we see that as force increases to 10 pN, the hinged-hinged and clamped-clamped results are close to each other. This is because the filament is stretched straight under large tension so that the tangents are aligned with the **e**_1_-axis even when the ends are hinged. Note that for the cantilever boundary condition, we observe significant curvature changes near the *s* = *L/*2 end for weak imposed tension but large activity. In general. the mean square displacements decrease as applied tension increases, and increase as active noise increases. However, the effect of increasing force is more significant than increasing active noise in the parametric range studied here. The result in Fig. 3 under small applied tension (*F* = 0.1pN) is in good agreement with the one from^35^.

We choose *L*/*ε* = 4 in studying the force-extension relations at three different activity values 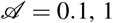, and 10 because entropic effects dominate for contour lengths which are long compared to the persistence length. Figure 3 shows our theoretical results; we see that under all magnitudes of activity, the force *F* needed to attain the same value of 〈*X*〉/*L* is the highest for the cantilever and the lowest for the clampedclamped condition. As 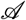 increases, the differences in the force-extension curves between these boundary conditions becomes smaller. Here the results have been plotted using *N* = 100 terms in the eqn. (58)&(59). To check convergence and accuracy, we estimated values of 〈*X*〉/*L* for *F* = 10pN and 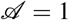. Summing *n* =10, 20, 50, 90, and 100 terms we find values 〈*X*〉/*L* = 0.7696, 0.7496, 0.7374, 0.7338 and 0.7333, confirming moderately rapid convergence and high accuracy.

### B. Activity from experimental measurements

Experiments of fluctuating filaments in active media form an integral part of the biophysical characterization of active or biological materials. In many of these cases, the activity of the ambient medium may not be known quantitatively. Here we discuss several ways of measuring the activity using expressions derived above. In each case it is assumed that the thermal temperature of the bath T, the contour length of the polymer *L*, the force exerted *F* are known. Properties of the active agents and the distribution of active sites encoded in *ℓ* is also assumed to be known along with drag and diffusion coefficients. For the specific case of a polymer in an active bath, these can be easily estimated from experiments without the polymer by visually tracking the self-propelling agents and computing correlation functions.

First, the effective bending modulus 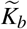 could be measured from a read-out of persistence lengths. We point out that eqn. (41) allows for extraction of information about the activity in a straightforward manner. Assume the experiment is conducted at large force, then from eqn. (29) we can derive a relation between the effective bending modulus 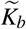, the purely thermal bending modulus *K_b_*, and *V*_0_:

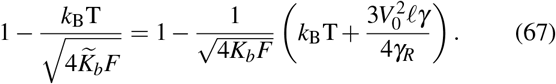

From a knowledge of 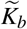, which is the apparent bending modulus in an active bath, and the original bending modulus *K_b_* we can then determine the velocity *V*_0_ representing the activity in the bath and/or filament.

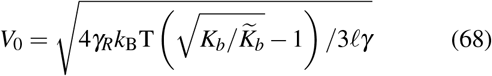

Note that use of eqn. (68) requires us to plug in values for the length scale *ℓ* and the friction coefficients *γ* and *γ_R_* which can be estimated by tagging the active agents/particles and measuring their single agent mean square displacements and re-orientation characteristics. This method is useful when the filament is long compared to its persistence length so that the end-to-end extension varies over broad range under various applied tensions.

In the case of clamped-clamped filaments one could measure the mean square deflection at the center *s* = 0 of the filament in an experiment to estimate *V*_0_ using

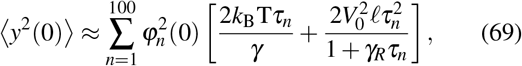

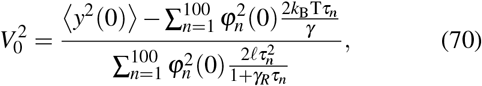

where eigenfunctions *φ_n_* are given in eqn. (53)&(55). Fig. 5(a) shows non-dimensional mean squared deflections plotted against 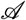 for various values of the tension. From a knowledge of mean square displacement one can read of the value of *V*_0_ using these curves. It may even be possible to measure 〈*y*^2^(*s*)〉 over the range −*L*/2 ≤ *s* ≤ *L*/2 for clamped-clamped or hinged-hinged boundary conditions with known applied forces on filaments and known bending moduli as in Arsenault *et al*^44^ (who applied forces using optical traps on actin filaments) and then fit the appropriate expressions with *V*_0_ as fitting parameter. This method is useful when the lengths of the filaments are comparable to their persistence length.

**FIG. 5.**
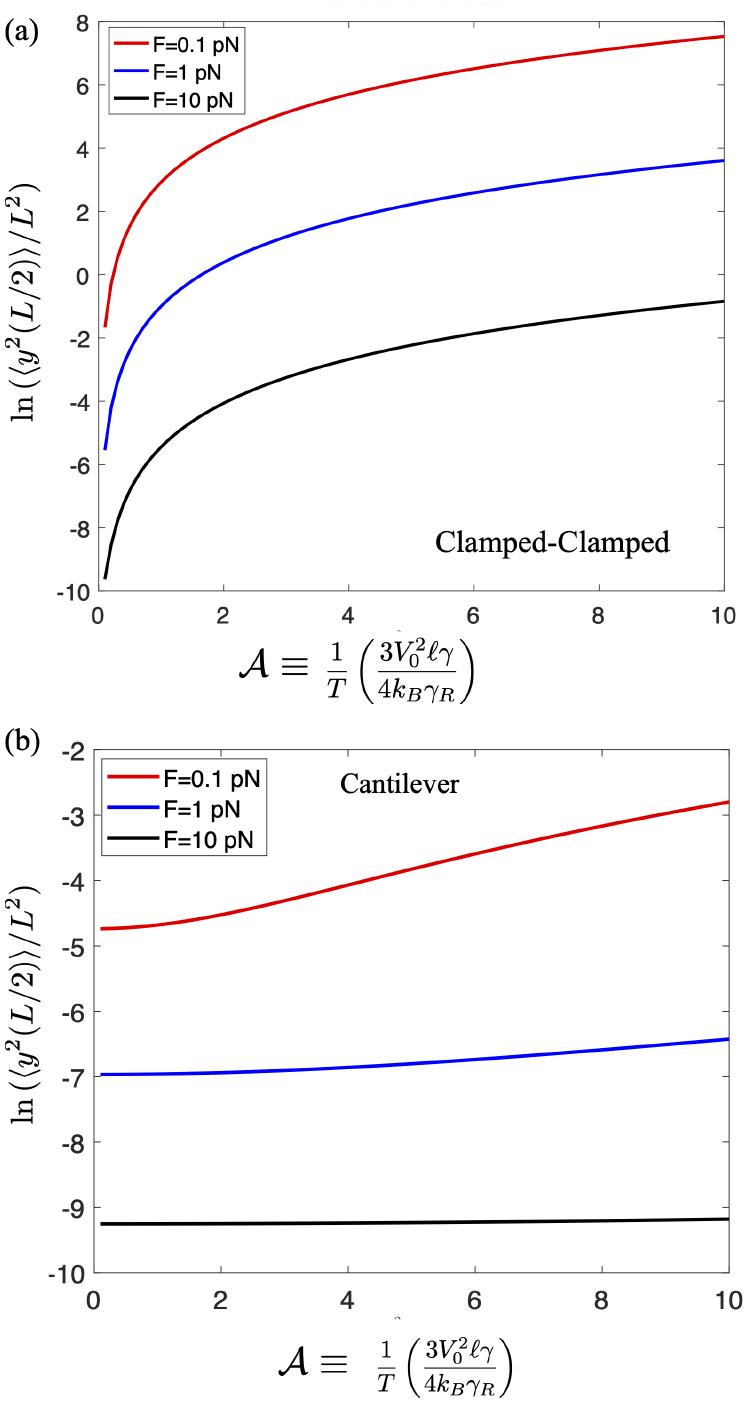
(a) Dimensionless mean square displacement at *s* = 0 as a function of dimensionless propulsion velocity under different applied tension *F* = 0.1 pN, *F* = 1 pN, *F* = 10 pN for a clamped-clamped filament. (b) Dimensionless mean square displacement at *s* = *L*/2 as a function of dimensionless propulsion velocity under different applied tension *F* = 0.1 pN, *F* = 1 pN, *F* = 10 pN for a cantilevered filament. Other parameters are the same as in Fig. 3.

In the case of cantilevered filaments (as in tethered particle experiments) one could measure the mean square deflection at the right end 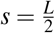 of the filament to estimate *V*_0_ using

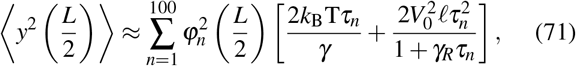

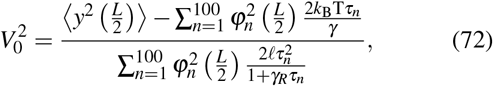

where eigenfunctions *φ_n_* are given in eqn. (66). Fig. 5(b) shows that the mean square deflection at the right end increases as the active agents increase under three different applied forces. However, this variation is not significant at large force (10pN).

## V. SUMMARY AND CONCLUSIONS

Filaments in live cells are subject to activity in addition to thermal Brownian motion due to the presence of molecular motors that when attached exert forces on these filaments. Often these active filaments and similarly activated filaments are subject to varying boundary conditions since they interact through cross-links or have other constraints placed on them^45,46^. Active filaments can be also synthetically manufactured by connecting polar and self-propelling colloids. In both these, and more general cases, the stochastic noisy nature of filament configurations arises from thermal noise and from athermal noise - the latter may originate from the interaction of the polymer with surrounding active (Brownian) agents or from the inherent motion of the polymer itself, which may be composed of active agents^47–50^.

Here, starting from a description of an active noisy filament as a continuous Gaussian semi-flexible polymer, we study how it responds under tension when constrained. Specifically, we derive the force-extension relations for the filaments under three distinct boundary conditions - hinged-hinged, clamped-clamped and a cantilever. We derive analytical expressions for the mean squared deflections and end-to-end distance (or mean stretch) of the polymer and study the variation of these with respect to activity and the magnitude of the applied force. The expressions for these quantities under hinged-hinged boundary conditions are reminiscent of the worm-like-chain model with a modified temperature that depends on the characteristic velocity of the activity. Our results provide methods to estimate the activity (of filaments or bath) by measurements of the force-extension relation of the filaments or their mean-square deflections which can be routinely performed using optical traps, tethered particle experiments, or other single molecule techniques.

Our approach can be extended to analyzing other problems that probe filament behavior in active suspensions. Recent analysis of constrained active filaments animated by follower forces^51,52^ such as induced by perfectly aligned molecular motors in microtubule and actin assays can be studied using an adaptation of eqn. (4). Similarly, the fluctuations of a spherical bead tethered to a surface by a slender semi-flexible polymer embedded in an active fluid comprised of self-propelling agents such as bacterial swarms, or photosensitive or phoretic colloidal suspensions can also be used to quantify bacterial activity. Using a bead that is ~ 10 – 20μm allows one to sample not just individual bacteria but collective velocity fields such as fluctuating vortices^53^. The force exerted by the diffusing bead *F* on the filament is here a fluctuating force that varies on time scales controlled by collective flow and agent-bead interactions.

## ACKNOWLEDGEMENTS

XL and PKP acknowledge support for this work through an NSF grant NSF-CMMI-1662101.

## Appendix A: Hinged-Hinged

Combining boundary conditions eqn. (30) with eqn. (12), we get four constraint equations,

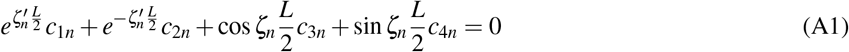

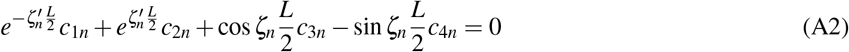

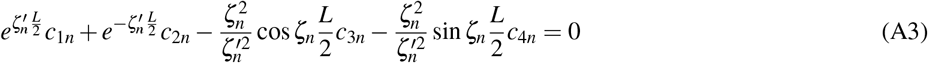

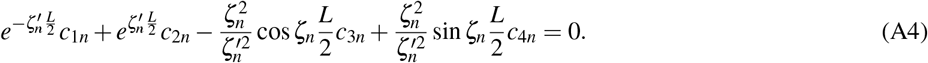

Comparing the expressions eqn. (A1)–eqn. (A3) and eqn. (A2)–eqn. (A4) (here ‘-’ means subtract), we deduce that

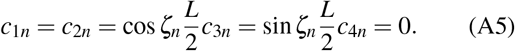

This indicates two sets of solutions,

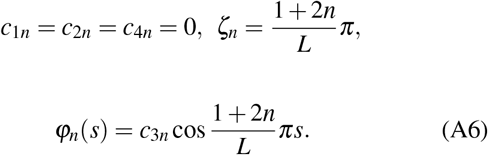

and

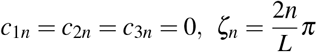

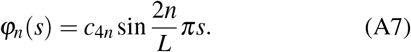

The general solution of *φ_n_*(*s*) could be obtained immediately through the combination of eqn. (A6) and eqn. (A8) as well as the constraint 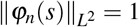,

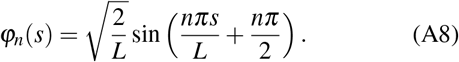

The first term in eqn. (33) can be summed as in^35^, and the result is,

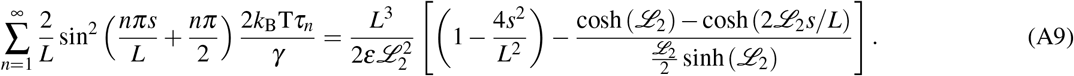

In order to compute the second term in eqn. (33), we introduce for brevity

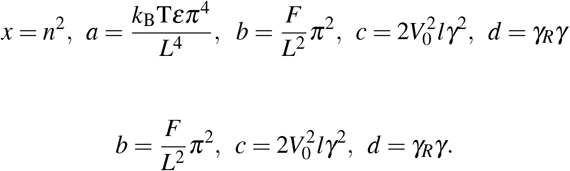

Using *x*_1_ and *x*_2_ as defined thus, we obtain

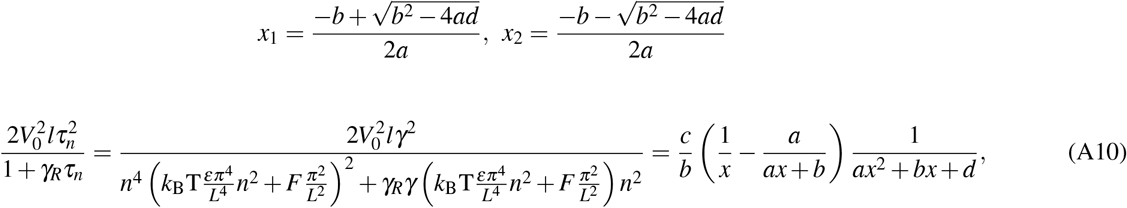

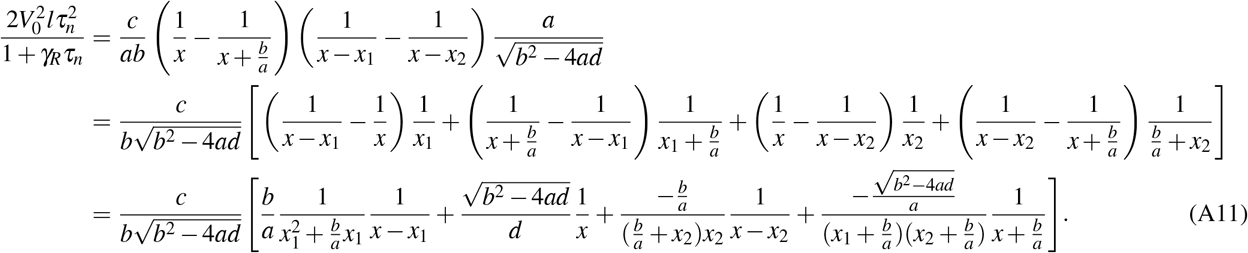

The second term in eqn. (33) can be simplified further using

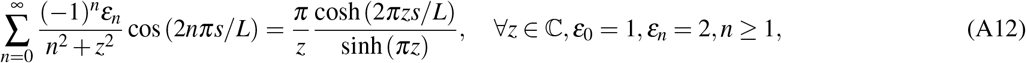

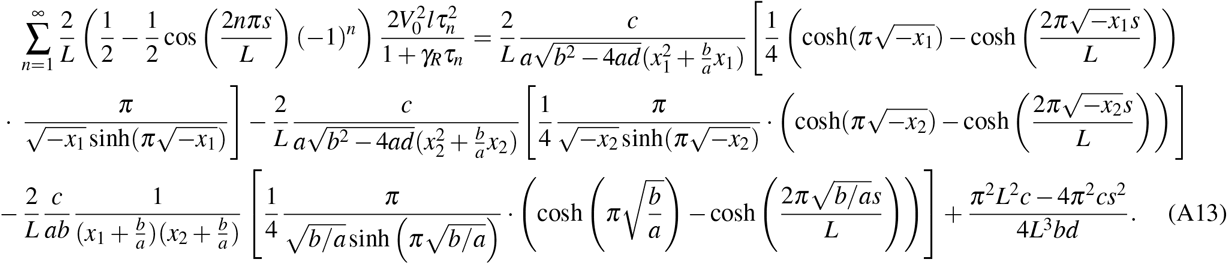

Setting *r*_1_ = −*x*_1_, *r*_2_ = −*x*_2_, then eqn. (26) can be summed to obtain eqn. (36). Let 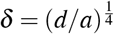. For the force extension relation, in the limit of 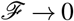, we obtain

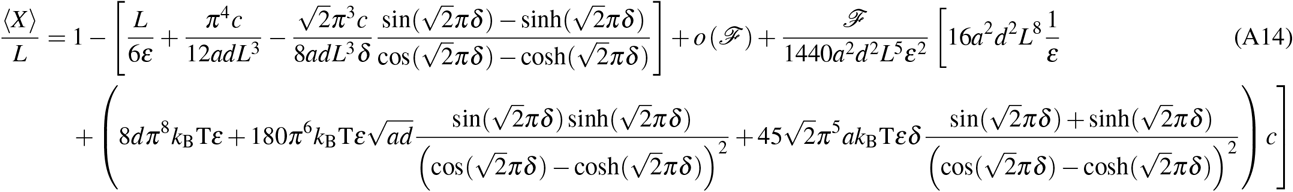

## Appendix B: Clamped-Clamped

After dragging eqn. (51) into eqn. (15), we have,

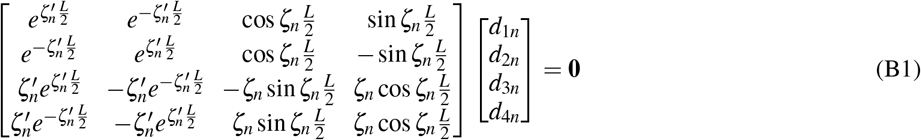

Using the Gaussian elimination, the coefficient matrix in eqn. (B1) can be simplified to,

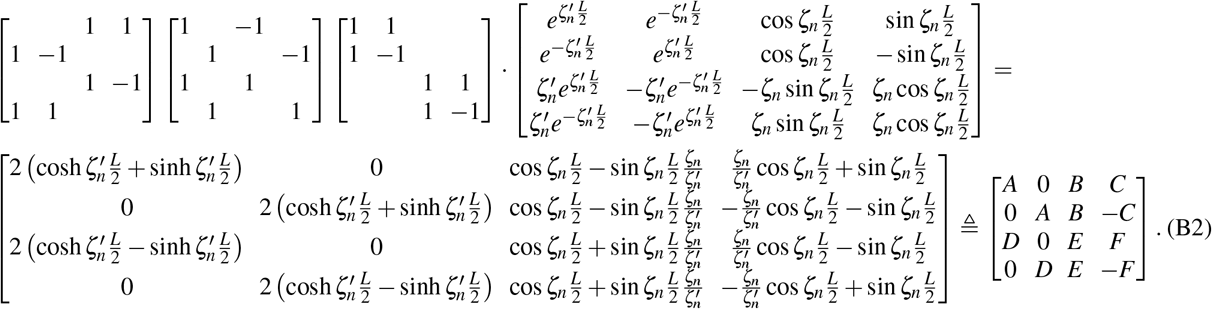

Applying Gaussian elimination once more on eqn. (B2), we get

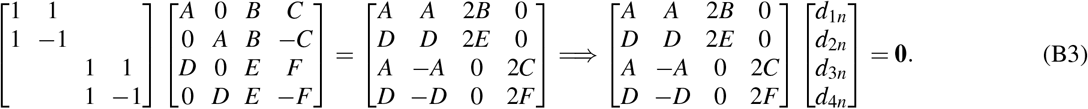

It clearly follows from eqn. (B3) that in order to derive non-zero solutions the only two possible cases are *AE* = *BD, AF* ≠ *CD* and *AE* ≠ *BD, AF* = *CD*. To see this, note that if *AE* ≠ *BD, AF* ≠ *CD*, then *d*_1*n*_ + *d*_2*n*_ = *d*_3*n*_ = *d*_1*n*_ – *d*_2*n*_ = *d*_4*n*_ = 0, ⇒ *d*_1*n*_ = *d*_2*n*_ = *d*_3*n*_ = *d*_4*n*_ = 0. Contradiction. if *AE* = *BD, AF* = *CD*, we’ll get 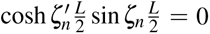 & 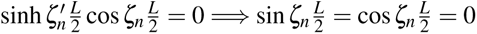. Contradiction. Now, if *AE* = *BD, AF* ≠ *CD*, then *d*_1*n*_ = *d*_2*n*_, 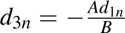, *d*_4*n*_ = 0, and we get eqn. (53). Similarly if *AE* ≠ *BD, AF* = *CD*, then *d*_1*n*_ = −*d*_2*n*_, *d*_3*n*_ = 0, *d*_4*n*_ = −*Ad*_1*n*_/*C*, and we get eqn. (55).

